# Correlated Protein Modules Revealing Functional Coordination of Interacting Proteins are Detected by Single-cell Proteomics

**DOI:** 10.1101/2022.12.18.520903

**Authors:** Mo Hu, Yutong Zhang, Yuan Yuan, Wenping Ma, Yinghui Zheng, Qingqing Gu, X. Sunney Xie

**Author notes:** M.H. and Y.Z. contributed equally to this paper.

## Abstract

Single-cell proteomics has attracted a lot of attention in recent years because it offers more functional relevance than single-cell transcriptomics. However, most work to date focused on cell typing, which has been widely accomplished by single-cell transcriptomics. Here we report the use of single-cell proteomics to measure the correlations between the translational levels of any pair of proteins in a single mammalian cell. In measuring pairwise correlations among ∼1,000 proteins in a population of homogeneous K562 cells in a steady-state condition, we observed multiple correlated protein modules (CPMs), each containing a group of highly positively correlated proteins that are functionally interacting and collectively involved in certain biological functions, such as protein synthesis and oxidative phosphorylation. Some CPMs are shared across different cell types while others are cell-type specific. Widely studied in omics analyses, pairwise correlations are often measured by introducing perturbations to bulk samples. However, some correlations of gene or protein expression in steady-state condition would be masked by perturbation. The single-cell correlations probed in our experiment reflect intrinsic steady-state fluctuations in the absence of perturbation. We note that observed correlations between proteins are experimentally more distinct and functionally more relevant than those between corresponding mRNAs measured in single-cell transcriptomics. By virtue of single-cell proteomics, functional coordination of proteins is manifested through CPMs.

**Table of Contents Image:** 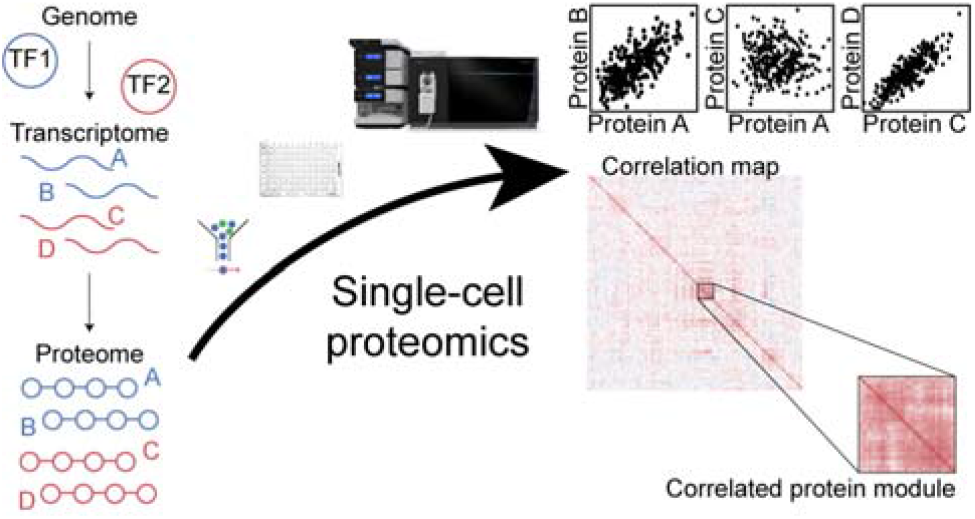

## Introduction

In a living cell, abundances of proteins and mRNAs fluctuate stochastically over time or even oscillate periodically (1, 2), hence cell-to-cell variations. This is because in a single cell there are only one or two copies of DNA (2–4), whose transcription and translation cannot be synchronized among different cells. On the other hand, a particular cellular function, such as protein synthesis, often requires multiple proteins, which, if each fluctuating, need to be synchronized in order for them to work together. While bulk measurement fails to study the ubiquitous fluctuation, variation of mRNA abundances and protein abundances can be revealed at single-cell level. Single-cell live imaging was able to detect synchronized fluctuations of functionally related molecules (5), however, it does not allow genome-wide measurement. Thanks to the single-cell omics technologies, the intrinsic fluctuation of gene expression, though difficult to probe in real-time, may be deduced through correlation analyses on single cells under the steady-state condition (6) (**Fig. 1a**).

**Figure 1.**
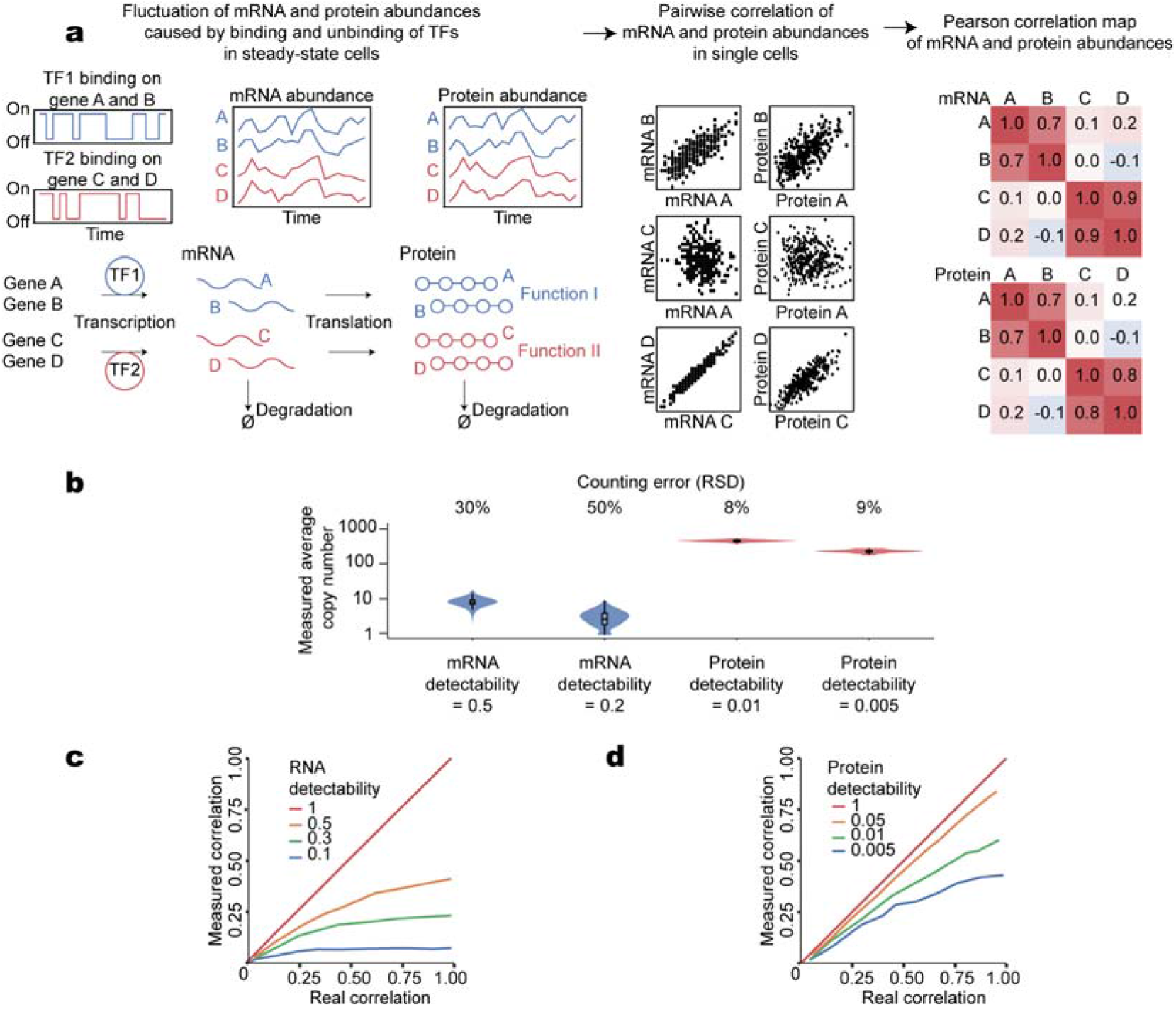
Direct measurement of protein correlations at single-cell level reveals intrinsic dynamics of the cell. a. Measurement of correlations of protein abundances in single cells reveals coregulation in steady-state gene expression. Levels of mRNAs and proteins fluctuate stochastically in steady-state cells, mostly caused by fluctuation in transcription factors (TFs). The expression of genes with similar biological functions tend to be coregulated, resulting in highly positive correlations of the abundances of corresponding mRNAs and proteins. Independently regulated proteins, on the other hand, will not exhibit high correlations. These correlations can be assessed by measuring abundances of mRNAs and proteins in single cells. Levels of mRNAs and proteins and their pairwise correlations are calculated from simulated theoretical distributions without considering detectability. b. The measured copy numbers and counting errors are affected by real copy numbers of protein/mRNA and detectability. c. Simulation of the effect of detectability on the measured correlation and real correlation in single cells for mRNA. d. Simulation of the effect of detectability on the measured correlation and real correlation in single cells for protein.

Currently, single-cell transcriptomics study has been widely used to distinguish different types of cells. Apart from cell typing, high-precision single-cell transcriptomics can also measure correlations in steady-state gene expression (6, 7). However, proteins, rather than mRNAs, are direct executers of most biological processes. Since protein abundances cannot be accurately predicted by mRNA levels (8–10), the dynamics of protein abundances and their correlations can only be assessed at the protein level. Moreover, small copy numbers of mRNA in single cells combined with limited detectability of current technologies (3, 11) would lead to high measurement noise (**Fig. 1b**), hindering accurate correlation measurements (**Fig. 1c**). Despite a much lower level of detectability and limited number of cells, correlations measured at the protein level proved to be more faithful than those measured at RNA level (**Fig. 1c, d**).

To investigate the functional correlations among proteins, various methods have been developed, including protein correlation profiling (PCP) (12–15), which is based on the hypothesis that interacting proteins (or proteins in the same pathway) should be positively correlated. Also, correlation-centric approach, with accurate quantification by mass spectrometry, proved to be powerful in capturing indirect interactions and proteins that interact but do not co-localize (16). However, these methods all rely on perturbations and bulk measurement where protein abundances are averaged across different cells, and cannot probe intrinsic dynamics of steady-state cells. Therefore, system-wide single-cell proteomics is needed.

Current mass spectrometry-based single-cell proteomics studies can detect and quantify hundreds or even thousands of proteins in a single cell using isobaric labeling (17, 18) or label-free (19, 20) quantification. Most existing studies focus on distinguishing different types of cells (17–19), which has been widely accomplished by single-cell transcriptomics analysis. Protein correlations of parts of the proteome are analyzed by single-cell proteomics (17, 21). However, to the best of our knowledge, no obseravation of protein modules by single-cell proteomics has been reported. To date, single-cell proteomics has not been used to study proteome-wide pairwise protein correlations of steady-state cells.

Here we report a robust mass spectrometry-based single-cell proteomic pipeline, which allowed us to measure correlations among ∼1,000 proteins in K562 cells. By measuring pairwise correlations based on protein abundance fluctuations in homogeneous cells, we identified correlated protein modules (CPMs), which are groups of proteins with highly positive pairwise correlations. CPMs were characterized by enrichment of certain cellular functions and known protein complexes. Furthermore, we demonstrated that CPMs identified from single-cell proteomic measurement could provide clearer and more solid evidence on functional relationships among proteins than single-cell RNA-seq based approach.

### Experimental and Theoretical Methods

#### Preparation of single-cell samples

K562 cells were resuspended from culture dish by cold PBS. 293T cells were treated by trypsin for 3 min before washed from dish by cold PBS. Then both cells were washed two more times by cold PBS. Single cells or 10 cells were sorted to 96-well plates (0030129512, Eppendorf) containing 0.5 µL 1% DDM in water by FACS (FACSAria, BD). Then plates were stored at -80 °C before use. Oocyte-cumulus complexes were collected from C57/6J strain mice superovulated with pregnant mare serum gonadotropin (PMSG) and HCG injections. Hyaluronidase was used to remove cumulus cells. MII oocytes were exposed to PBS solution. Single oocyte cells were then picked by pipetting with an aspirator tube to 0.2 mL PCR tubes (PCR-02-L-C, Axygen). Then cells were stored at -80 °C before use.

### Proteomic sample preparation

We used an acoustic liquid handling workstation (Echo 525, Beckman) to increase throughput and process small volumes of liquids. Sorted single cells in 96-well plate or 200 µL PCR tubes were digested directly by trypsin at 37 °C for 3 hours. For label-free pipeline, 4.4 µL 0.43% TFA and 1% acetonitrile in water was added to terminate digestion (3.5 µL for oocytes). Then peptides were dried in a concentrator (LabCondo) at 45 °C. 5 µL of 0.1% TFA and 1% ACN in water were used to resuspend the peptides. Digests were transferred to sample tubes (186009186, Waters) for LC-MS/MS.

### LC-MS/MS

4 µL of the digests were injected. Peptides were separated by a commercialized high-performance chromatography column (AUR2-25075C18, IonOpticks) at 100 nL/min on a nanoflow liquid chromatography (Ultimate 3000 RSLCnano, ThermoFisher). For label-free pipeline, the effective gradient was 70 min, generating throughput of 16 cells per day (full gradient of 90 min). Details of gradient are described in the Supporting Information.

Peptides were analyzed by a tribrid mass spectrometer (Orbitrap Eclipse, ThermoFisher) with an ion mobility interface (FAIMS Pro, ThermoFisher). FAIMS compensation voltages of -55 V and -70 V were used. Cycle time of 1 s was used for both CVs. MS spectra were acquired by Orbitrap analyzer, and MS/MS spectra were acquired by linear ion trap analyzer. Max ion injection time for MS/MS was 200 ms.

### Proteomic data analysis

MS raw files were searched against UniProt human protein database (20,286 reviewed accessions, downloaded on April 14, 2020, for searches of mouse oocytes, a FASTA database with 17,015 reviewed accessions downloaded on July 31, 2020 were used instead) and an in-house curated contamination database (284 accessions) by Proteome Discoverer 2.4 (ThermoFisher). For identification, 1% FDR for PSMs, peptides and proteins were used. For label-free quantification, peak intensity of MS feature was used as signal of quantification and match-between-runs (MBR) feature was used. Detailed data analysis method of Proteome Discoverer can be found in data repository (https://doi.org/doi:10.25345/C55717S21).

### Generation of correlated protein modules

The abundances of all proteins in quantified cells were further filtered to 1) remove cells with extremely high level of contamination; 2) remove protein contaminants; 3) remove proteins with more than 50% missing values in all cells; 4) remove outlier cells (by fitting to a Gaussian distribution). Protein abundance of single cells was normalized by total protein intensity. The missing values were filled with zeros. Pearson correlation of proteins were calculated by numpy.corrcoef of NumPy (22). The correlation map was reordered using unweighted pair group method with arithmetic means clustering using the SciPy (23) package. Llists of correlation coefficient of proteins in K562, 293T and mouse oocyte can be found in the Supporting Information, Dataset S1-S3. To obtain a CPM within a correlation map, first, protein groups that have over 80 percent of protein pairs with correlation coefficient over 0.3 are identified. Then, among these protein groups, the one with the largest number of proteins is defined as a CPM. A list of correlated protein modules can be found in the Supporting Information, Dataset S4. We define the abundance of a CPM in a cell as the sum of the abundances of all proteins included in the very CPM.

### Bioinformatic analysis

Bioinformatical analysis was conducted with R. Principle component analysis was conducted by “PCA” function in “FactoMineR” package with default parameters. Gene Ontology enrichment and KEGG pathway enrichment analysis was done by R package “clusterProfiler” (v.4.0.2). The adjusted p-value cutoff was set as 0.05, and only GO terms with more than 2 genes were kept for further analysis. Reactome pathway enrichment analysis was done by R package “ReactomePA”. The p-value cutoff was set as 0.05, and only terms with over 2 genes were kept. Protein-protein interaction ratio was calculated based on R package “STRINGdb” (v.11). The combined score cutoff was 250. The ratio value was calculated as the proportion of known PPI pairs to the total number of protein pairs in a CPM. CORUM version 3.0 dataset was downloaded at “http://mips.helmholtz-muenchen.de/corum/“. Results of enrichment analysis of correlated protein modules can be found in the Supporting Information, Dataset S4. To assess the statistical significance of the CPM k with N proteins, N proteins were randomly selected from the correlation map for 10,000 times. For the i^th^ group of selected proteins, the average correlation is Cor_i_. The average correlation of the CPM k is Cor_k_. The bootstrap P value is given by:

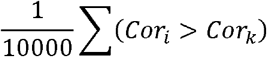

The P values are adjusted by Benjamini-Hochberg method with p.adjust() function in R package “stats”.

### Simulation of effect of detectability on measured correlation and real correlation

To assess the measurement noise of single-cell RNA analysis and single-cell proteins analysis, simulation was conducted with R. Random RNA abundances of 1000 cells were generated by Poisson distributions with median of 17 copies. Random protein abundances of 100 cells were generated by negative bimodal distributions with mean value of 5*10^4^ and CV of 0.01. The measured copy number was obtained after multiplying detectability by RNA/protein abundances. Finally, the measurement noise was calculated as relative standard deviation (RSD) of measured copy number and real (randomly generated) copy number.

To assess the accuracy of correlation measurement of RNA and proteins, simulation was conducted with R. Random RNA abundances of 1000 cells were generated by Poisson distributions with median of 20 copies. Random protein abundances of 100 cells were generated by negative bimodal distributions with median of 10^5^ and CV of 0.01. Then a normally distributed random noise was added to RNA and protein. Next, measured copy number was obtained after multiplying detectability by RNA/protein abundances. Finally, the accuracy of correlation measurement was calculated by comparing measured copy number with real (randomly generated) RNA/protein abundances. All simulations were repeated 100 times to reduce random noise.

### Comparison of CPMs with CGMs and ProteomeHD

The correlation coefficient and CGMs of 12,660 genes in 2,302 K562 cells are analyzed as described in (6). When comparing CPMs and CGMs, because there are multiple transcripts corresponding to the same protein, the number of proteins and genes in **Fig. 3a** are not the same. The correlation scores of ProteomeHD (17) are no more than 0.3. To compare with our Pearson correlation coefficient, ProteomeHD data was scaled by making its maximum correlation score equal to 1.

## Data and code repositories

The proteomic datasets can be found at MassIVE (https://doi.org/doi:10.25345/C55717S21). The code for processing data can be found at GitHub (https://github.com/dionezhang/CPM).

## Results and Discussion

### Correlated protein modules in K562 cell line

To measure pairwise correlations between proteins, protein abundances need to be measured accurately in the first place. To this end, we optimized the label-free single-cell proteomic pipeline (**Fig. 2a**) by balancing the throughput, coverage of proteome and robustness of the quantification performance. Longer gradient generally improves the protein identification, but reduces the number of cells to be analyzed each day. Therefore, we ran single-shot LC-MS experiments at a moderate gradient length (16 cells per day).

**Figure 2.**
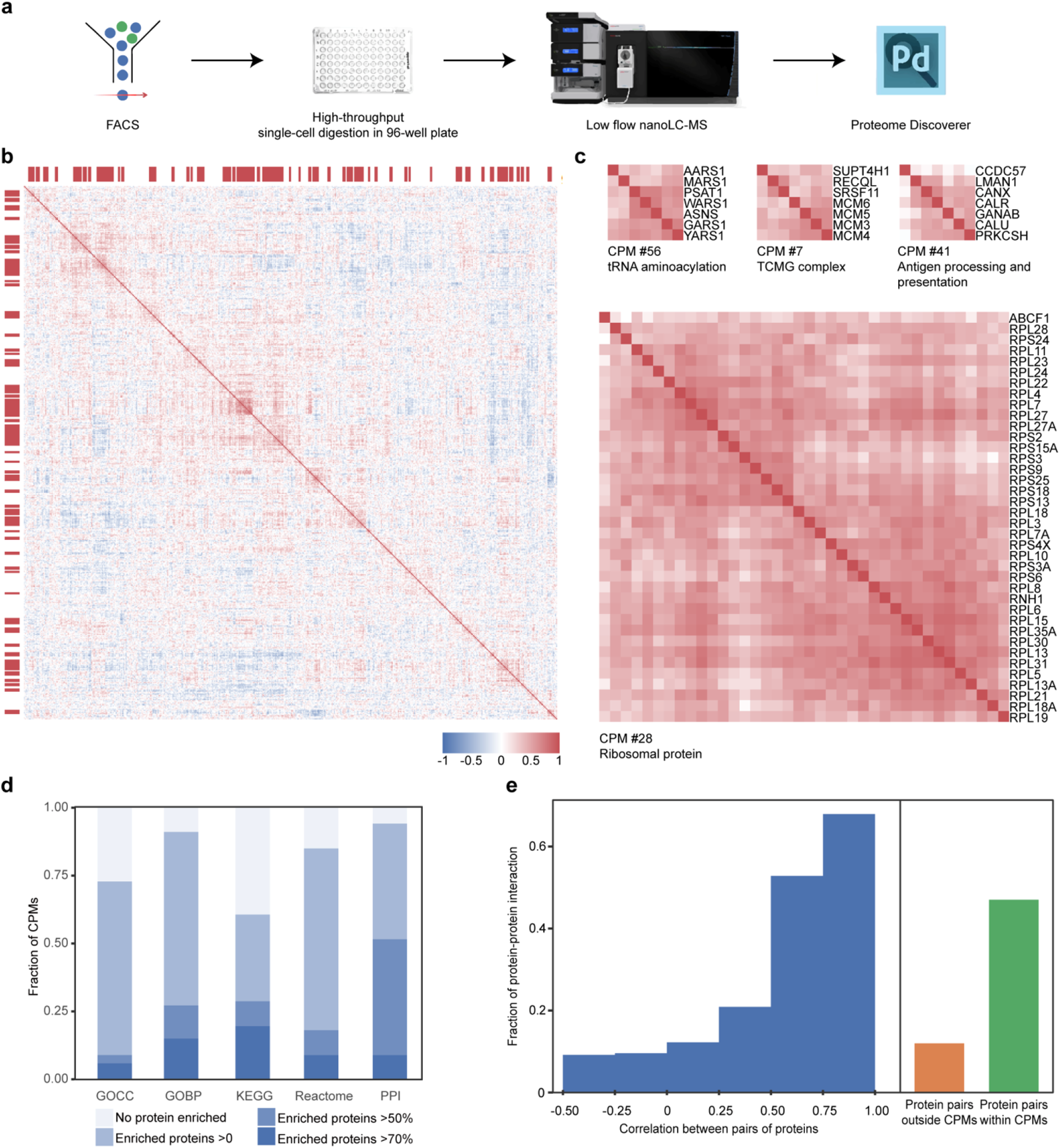
Correlated protein modules of K562 cell line by single-cell proteomics. a. Flowchart of pipeline for measuring protein-protein correlations in single-cells. b. Correlation map of 1,249 proteins in 69 K562 single cells. Red blocks indicate that corresponding proteins were defined as a CPM. c. Four representative CPMs and their enriched function terms. d. Fraction of CPMs with different enrichment results of Gene Ontology cellular component (GOCC), Gene Ontology biological processes (GOBP), KEGG, Reactome and protein-protein interaction (PPI) of proteins in each CPM. CPMs were categorized by the ratio of proteins with enriched functional terms (enriched proteins) to total proteins in the corresponding CPM. e. Fraction of protein pairs with known PPIs in the STRING database at different levels of correlation coefficients (blue), outside CPMs (orange), and within CPMs (green).

To test the performance of our pipeline, we analyzed single K562 cells and single 293T cells. After removing contaminations, low-quality proteins and ourlier cells, our pipeline clearly distinguished between the two cell populations (**Fig. S1a**). This encouraged us to further test its ability to measure protein-protein correlations in a homogeneous cell line. In 69 K562 cells, 1,249 proteins were quantified. Pearson correlation coefficients of the abundances of these proteins centered around zero (**Fig. S1b**). After unsupervised hierarchy clustering based on Pearson correlation coefficients of protein abundances, we found groups of highly positively correlated proteins, named as correlated protein modules (CPMs). 66 CPMs were found in K562 cells, containing 546 proteins in total (**Fig. 2b**). Each CPM contained 5 to 38 proteins. The correlation coefficients between protein pairs within a CPM, with average values ranging from 0.29 to 0.53, were much higher than those outside CPMs (**Fig. S2a**).

To examine potential functional relationships among proteins in each CPM, we compared our data with CORUM (24) to see if CPMs represent certain protein complexes. Several subunits from the same complex appeared in the same CPM. For example, CPM #7 contained 5 spliceosome subunits and CPM #34 contained 4 EIF subunits. CPM #19 was almost entirely composed of T-complex proteins (5/7) and CPM #28 mostly consisted of ribosomal proteins (37/38).

Proteins in the same CPMs tend to have the same subcellular localization, as shown by enrichment analysis of the Gene Ontology cellular component (GOCC). For example, CPM #15 was enriched with the ribonucleoprotein complex and telomeric region while CPM #7 was enriched with the CMG complex. Over half of the CPMs had significant enrichment of GOCC terms.

Functional relationships indicated by the Gene Ontology biological process (GOBP), KEGG pathway (25) and Reactome (26) pathway enrichment were highly prevalent among CPMs (**Fig. 2d, Fig. S3)**. For example, CPM #56 was enriched with “tRNA aminoacylation (GO:0006418)” while CPM #41 was enriched with “antigen processing and presentation (KEGG: hsa04612)”. That most CPMs were significantly enriched with GOBP terms suggests that involving in the same biological process is a major contributor to protein correlations in steady state.

Some CPMs were enriched with multiple functions. For example, in CPM #6, TRIM28 and USP7 were involved in “DNA demethylation (GO:0080111)” while DCD and S100A9 were related to “antimicrobial humoral response (GO:0019730)”. This implies that DNA demethylation and antimicrobial humoral response are closely related, consistent with a previous study (27).

Finally, we examined the relationship between each protein pair within a CPM in the STRING database (28) to see how well protein correlations reflect actual protein-protein interactions. It was found that all CPMs except four had corresponding PPIs. Over 90% of protein pairs in CPM #56 and CPM #61 had known PPIs in the STRING database. To further evaluate the relations between pairwise protein correlations, CPMs and protein-protein interactions, we calculated the fraction of known protein-protein interaction at different correlation level between pairs of proteins in steady-state single-cells. For a pair of proteins, a larger correlation coefficient means higher possibility that the two proteins have known interactions (**Fig. 2e**). We also found that protein pairs within CPMs are much more likely to have known PPI in the STRING database compared to a pair of proteins not in the same CPM (**Fig. 2e**), which is consistent with the fact that CPMs reflect functional coordination of interacting proteins.

As CPMs enrich specific cellular functions, correlations between CPMs may imply relationships between different cellular activities. We defined the abundance of a CPM as the sum of the abundances of all proteins included in that CPM. We further measured the correlation between CPMs by calculating the pairwise correlations of abundances of CPMs. Most of the CPMs were poorly correlated, with correlation coefficients from 0 to 0.2. However, some CPM pairs showed negative correlations lower than -0.3, which means the abundance of one CPM goes up in a certain cell when the other one goes down. For example, CPM #11 and CPM #57 had a negative correlation coefficient (**Fig. S2b**). The 11 proteins in CPM #11 were related to “rRNA processing (GO:0006364)” and “DNA post-replication repair (GO:0006301)”, while the 9 proteins in CPM #57 were involved in “ATP generation (GO:0006757)” and “NADH metabolism (GO:0006734)”. For cells that had high CPM #11 abundance, the abundance of CPM #57 was rather low, consistent with a previous study reporting that ribosome biosynthesis is controlled by fluctuation of NAD(+)/NADH ratio (29). CPM #57 was also negatively correlated with CPM #37 (**Fig. S2c)** which contains proteins TOP2A and CDK1. These two proteins were found to be involved in regulating the circadian rhythm (30). Thus, this negative correlation suggested that CPM #57 was a circadian rhythm-dependent CPM.

### CPM is a shared feature of functional proteome

To verify whether CPM is a shared feature of functional proteome, we further examined protein correlations in the human 293T cell line and mouse MII oocytes by single-cell proteomics. The correlation map of 1,150 proteins in 63 293T cells revealed 50 CPMs (**Fig. S4a**). Despite the differences between K562 and 293T cell line in protein levels and in sizes of CPMs, the functions of CPMs in two cell lines appeared similar. For example, T-complex CPM was also found in 293T cells (**Fig. S4d**) as in K562 cells. Proteins of tRNA aminoacylation module in K562 were also highly correlated in 293T, despite of marked differences between the two cell lines in the abundance of these proteins (**Fig. S4c**). This shows that although abundance of proteins with the same function may vary among different types of cells, the synchronized fluctuation of them is a consistent feature of functional proteome.

Experiments on 137 mouse MII oocytes quantified 3,422 proteins, which is about three-fold the number of quantified proteins in regular-size mammalian cells. We discovered 86 CPMs from these proteins (**Fig. S4b**). Compared with K562 and 293T, CPMs of mouse oocytes are generally bigger in size and tend to infer more complex biological functions. CPMs of certain protein complexes identified in K562 and 293T cells, such as T-complex (**Fig. S4d, e**), were also found in mouse oocytes, indicating that these CPMs are conserved across different species and cell types. CPMs associated with certain functions, such as proteasome complex, could only be identified in oocytes (**Fig. S4f**). Therefore, the modules identified from single-cells can infer cellular characteristics and are affected by the total number of identified proteins.

### CPMs reveal more functional information than CGMs and bulk proteomic approach

Similar to CPMs, correlated gene modules (CGMs) have also been reported based on pairwise correlations of RNA expression levels via MALBAC-dt, a single-cell RNA-seq method with high detectability and high accuracy (6). To compare the efficacy of CPM versus CGM in functional analyses, we examined the correlation coefficients of genes that have been identified from both methods. Transcripts of most identified proteins were found in MALBAC-dt data. Measured correlation coefficients of RNAs were more concentrated around zero compared to those of proteins (**Fig. 3a**). We also found the pairwise correlation coefficients of proteins in functional modules are generally higher than those of corresponding RNAs. For example, for the ribosome module, the average correlation coefficients among proteins is 0.55, compared with 0.32 for RNAs (**Fig. 3b**). Thus, compared to transcripts of ribosomal proteins, the correlation among ribosomal proteins is more distinguishable from the overall pairwise correlations (**Fig. 3c**). Furthermore, some functional modules identified at the protein level cannot be observed at the RNA level, such as CPM #11 (rRNA processing and mismatch repair) and CPM #3 (inner mitochondrial membrane oxidation phosphorylation) (**Fig. 3d**). These differences are likely due to the limited detectability of single-cell RNA-seq which masks the correlation of low-abundance transcripts, or to the possibility that some genes with a shared function are synchronized at protein level but not at transcription level. Therefore, CPM can provide more straightforward and relevant information on how genes work together to carry out biological functions.

**Figure 3.**
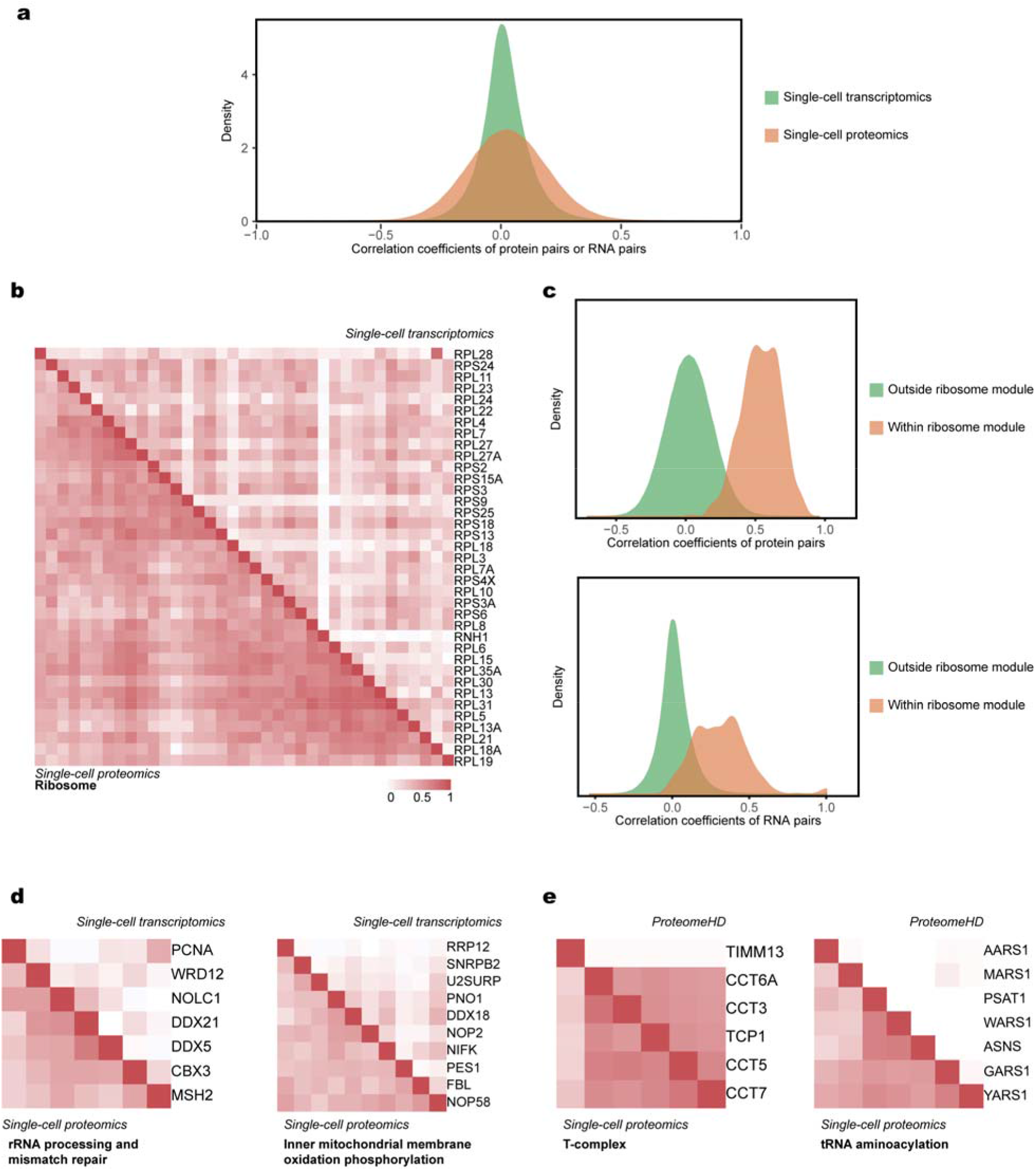
Correlated protein modules by single-cell proteomics exhibit unique characteristics. a. Distribution of pairwise correlation coefficients of single-cell transcriptomics (*N* = 1,275) and single-cell proteomics (*N* = 1,249). b. Comparison of protein (bottom-left) and mRNA (top-right) correlation coefficients for the ribosome modules of K562. Correlation coefficients between proteins are generally stronger than those between corresponding mRNAs. c. Distribution of correlation coefficients of protein pairs (top) or RNA pairs (bottom) within the ribosome module (*N* = 1,332 for proteins, *N* = 1,351 for RNAs) and outside the ribosome module (*N* = 1,558,752 for proteins, *N* = 1,624,981 for RNAs). The correlation coefficients of functional protein pairs are more distinguishable from the overall pairwise correlations. d. Comparison of protein (bottom-left) and mRNA (top-right) correlation coefficients for the module of rRNA processing and mismatch repairment and the module of inner mitochondrial membrane oxidation phosphorylation in K562. Correlation coefficients between proteins are generally stronger than those between corresponding mRNAs. e. Comparison of correlation coefficients in CPM (bottom-left) and scaled correlation scores from ProteomeHD (top-right) for the T-complex composition and tRNA aminoacylation modules. CPM clearly exhibits correlations that cannot be seen in the bulk data obtained via perturbation.

We then compared CPMs identified from our analysis with those discovered by bulk proteomics measurement under perturbations (16). Although some modules, such as ribosome and T-complex, were observed in both datasets, some biological processes like tRNA aminoacylation (CPM #56) were not found in bulk data (**Fig. 3e**). This shows that our method, which relies on single-cell measurement without perturbations, is more sensitive than bulk proteomics measurement in probing intrinsic correlations among proteins.

High-throughput single-cell analysis has proved to be a valuable strategy for understanding the functional mechanisms of cells and tissues (7, 31). However, identifying functional gene expression programs from large-scale single-cell dataset has been challenging. Most methods emphasize on heterogeneity among cells, such as principal component analysis (PCA) (32, 33) or non-negative matrix factorization (NMF) (34). These methods can uncover the variation of gene expression and cellular activity among different cell types, but cannot identify the networks of gene expression in homogeneous cell populations. Nonetheless, investigating correlations on the proteome level is still pertinent. This is because while mRNA fluctuation arises from stochastic expression of a single copy of gene in a live cell (2, 4), protein fluctuation arises from additional variation including translation, post-translational modification and protein degradation (9). Thus, direct measurement of protein correlation is required to uncover the intrinsic dynamics of protein abundance in steady state. Moreover, as the copy number of proteins in a single cell is much larger than that of mRNA, measurement of protein can reveal correlation of genes at much higher accuracy.

Here, through global correlation analysis of single-cell proteome measured by mass spectrometry, we revealed the functional coordination of proteins despite stochastic gene expression under steady-state conditions. Our method probed the intrinsic fluctuations of more than a thousand of proteins and identified groups of proteins that were highly correlated. Based on functional analysis of CPMs and the relationships between different CPMs, we discovered coordinated fluctuation for proteins involved in the same functions, related pathways or conflicting processes, which is inaccessible by other single-cell methods and bulk approaches.

The functional coordination and interaction of proteins is a shared feature among different types of cells, yet it is also determined by the cellular characteristic and affected by the detectability of the protein measuring technology. Currently, the proteome coverage of single-cell proteomics is still limited, especially when compared to single-cell transcriptomics which can identify more than 5,000 genes in a single cell. Therefore, while most key regulators in signaling networks, such as transcription factors, can be readily measured by current single-cell transcriptomics, these low-abundance proteins cannot be detected by single-cell proteomics yet.

In our work, we demonstate the use of label-free single-cell proteomics in the identification of correlated protein modules. However, our label-free quantification pipeline has limited proteome coverage and cell throughput (up to 40 cells per day (19, 35)). As the proteome coverage of single-cell proteomics improves, we may be able to gain more functional insights from CPM, such as coregulation mechanism of transcription factors. This possibility is showcased by our mouse oocyte dataset, where the total protein amount per cell is about 100-fold of that in K562. Therefore, substantial effort is needed to make CPM widely applicable to cell cultures or tissues, where much larger number of cells need to be measured and higher proteome coverage is preferred. This is possible given the rapid advances in sample preparation (36–38), LC separation (39, 40), MS instrumentation (35), data acquisition (41), and informatics (42, 43). We believe that CPM is a good demonstration of single-cell omics methods for functional annotation of the proteome.

## Conclusions

We have shown that accurate single-cell measurements of protein abundances unraveled the CPMs, which are otherwise masked by bulk MS measurements, and are more accurate than single-cell transcriptome measurements. A CPM is a group of proteins that collectively carries out specific biological functions, therefore their fluctuation must be correlated. Our results have demonstrated that single-cell proteomics can be used to probe protein-protein interactions and to study how proteins work together at a system-wide level.

### Supporting Information Description

Additional experimental details and data, including the full LC gradient of the label-free pipeline, four supporting figures (PDF), and supporting datasets (XLSX).

## Acknowledgement

This work was supported by Beijing Advanced Innovation Center for Genomics at Peking University and Beijing Municipal Commission of Science and Technology (Z201100005320015). We thank Alec R. Chapman, David F. Lee, Fuchou Tang and Ge Gao for their inspiring work in gene modules by high-precision single-cell transcriptomics. We thank Alec R. Chapman and Jiekun Yang for their involvement in the early attempt on single-cell proteomics of our group. We thank Bogdan Budnik and Nikolai Slavov for their help on our implementation of single-cell proteomics. We thank Xiaoran Chai and Wenjie Sun for helpful discussions of the manuscript.

