## supplemental text and figures for "Correlated Protein Modules Revealing Functional Coordination of Interacting Proteins are Detected by Single-cell Proteomics"

Mo Hu[abc§], Yutong Zhang[abd§], Yuan Yuan[abd], Wenping Ma[abe], Yinghui Zheng[ab], Qingqing Gu[c], X. Sunney Xie[abc*]

[a]Beijing Advanced Innovation Center for Genomics, Peking University, Beijing, 100871, China

[b]Biomedical Pioneering Innovation Center (BIOPIC), Peking University, Beijing, 100871, China

[c]Changping Laboratory, Beijing, 102206, China.

[d]Collage of Chemistry and Molecular Engineering, Peking University, Beijing, 100871, China

[e]Peking-Tsinghua Center for Life Sciences (CLS), Academy for Advanced Interdisciplinary Studies, Peking University, Beijing, 100871, China

[§]M.H. and Y.Z. contributed equally to this paper.

**This PDF file includes:**

Supporting information text
Figures S1 to S4

**Supporting Information Text**

Full LC gradient of the label-free pipeline

| Time | % B | Flow rate (nL/min) |
|---|---|---|
| 0 | 5 | 300 |
| 2 | 6.2 | 300 |
| 2.1 | 6.2 | 100 |
| 42 | 31.2 | 100 |
| 58 | 42.5 | 100 |
| 63 | 99 | 100 |
| 67 | 99 | 100 |
| 68 | 99 | 300 |
| 73 | 99 | 300 |
| 73.1 | 5 | 300 |
| 80 | 5 | 300 |

**Supporting Information Figures**

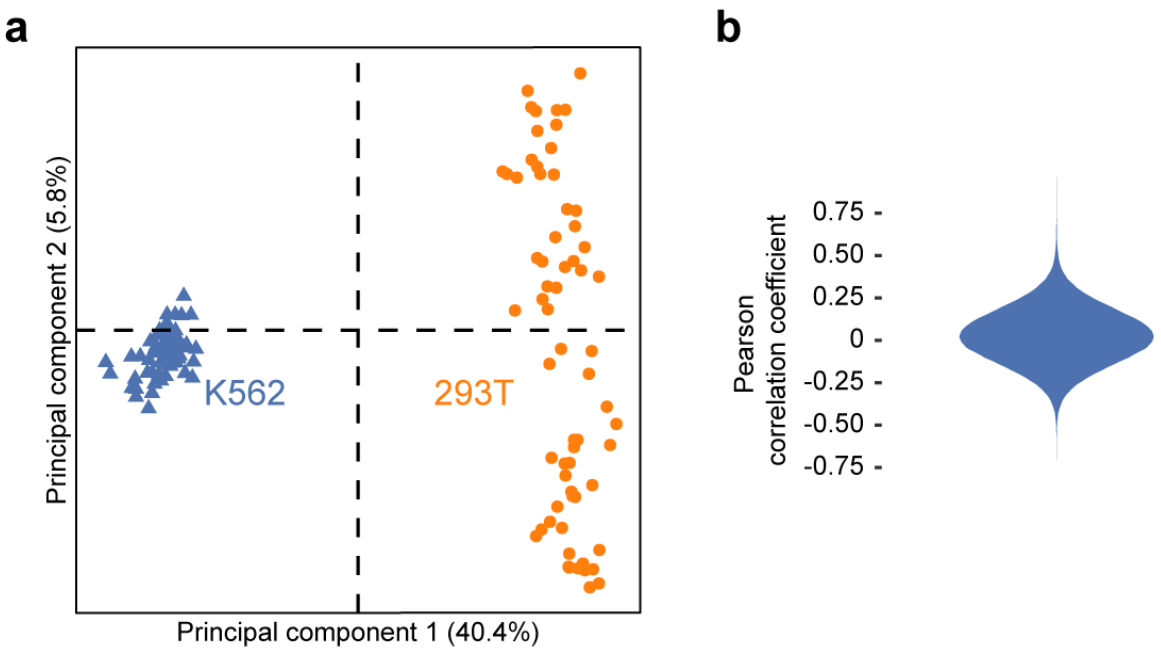


**Fig. S1.** Validation of the label-free single-cell proteomic pipeline. a. Principal component analysis of single K562 cells and single 293T cells. b. Violin plot of Pearson correlation coefficients of proteins of K562 ($N$ = 1,249).

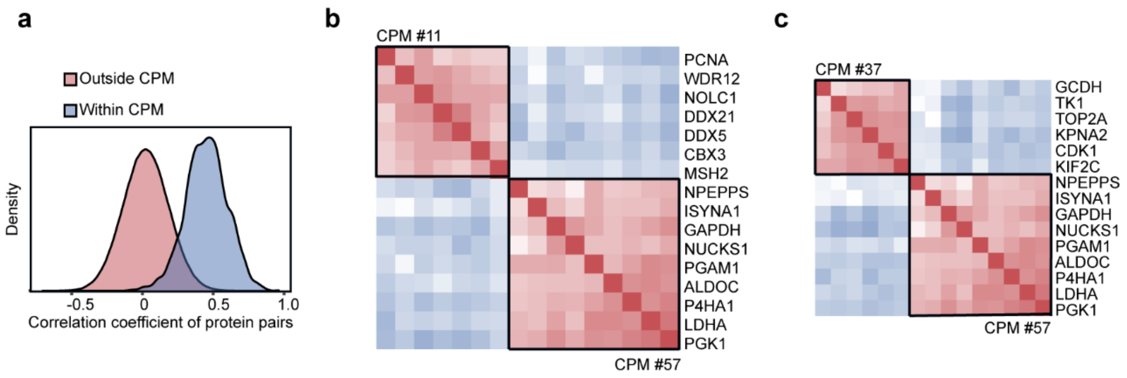


**Fig. S2.** Analysis of CPMs in K562. a. Distribution of Pearson correlation coefficients of protein pairs within CPM (blue, $N$ = 5,426) and outside CPM (red, $N$ = 1,553,326). b. Correlation between CPM #11 and CPM #57. The order of proteins was kept as the original order in correlation map. c. Correlation between CPM #37 and CPM #57. The order of proteins is kept as the original order in correlation map.

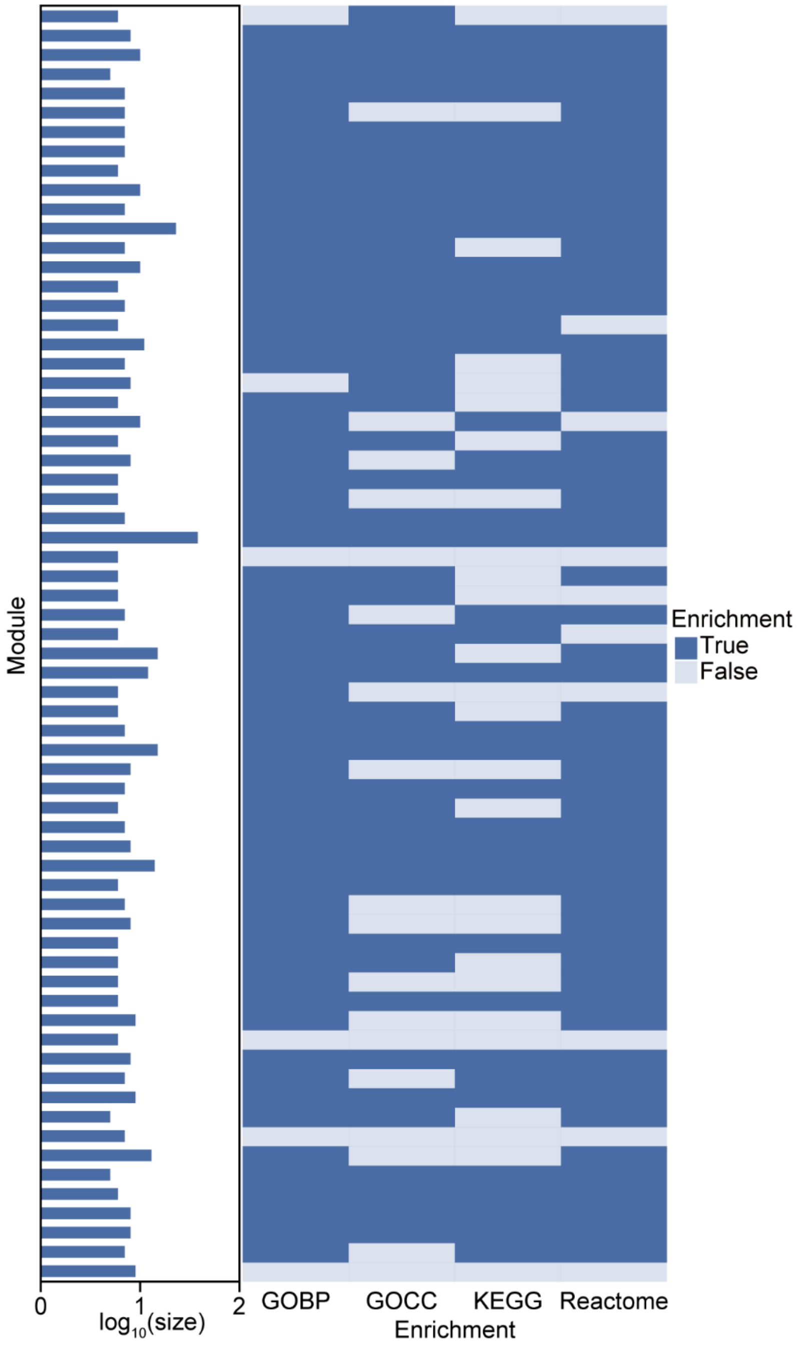


**Fig. S3.** Enrichment analysis of CPMs in K562. Each row is a CPM from K562 dataset. Left: the bar plot shows the size of each CPM, which is the number of proteins in each module. Right: the heatmap shows the enrichment results of Gene Ontology cellular component (GOCC), Gene Ontology biological processes (GOBP), KEGG pathways, and Reactome pathways (FDR < 0.05).

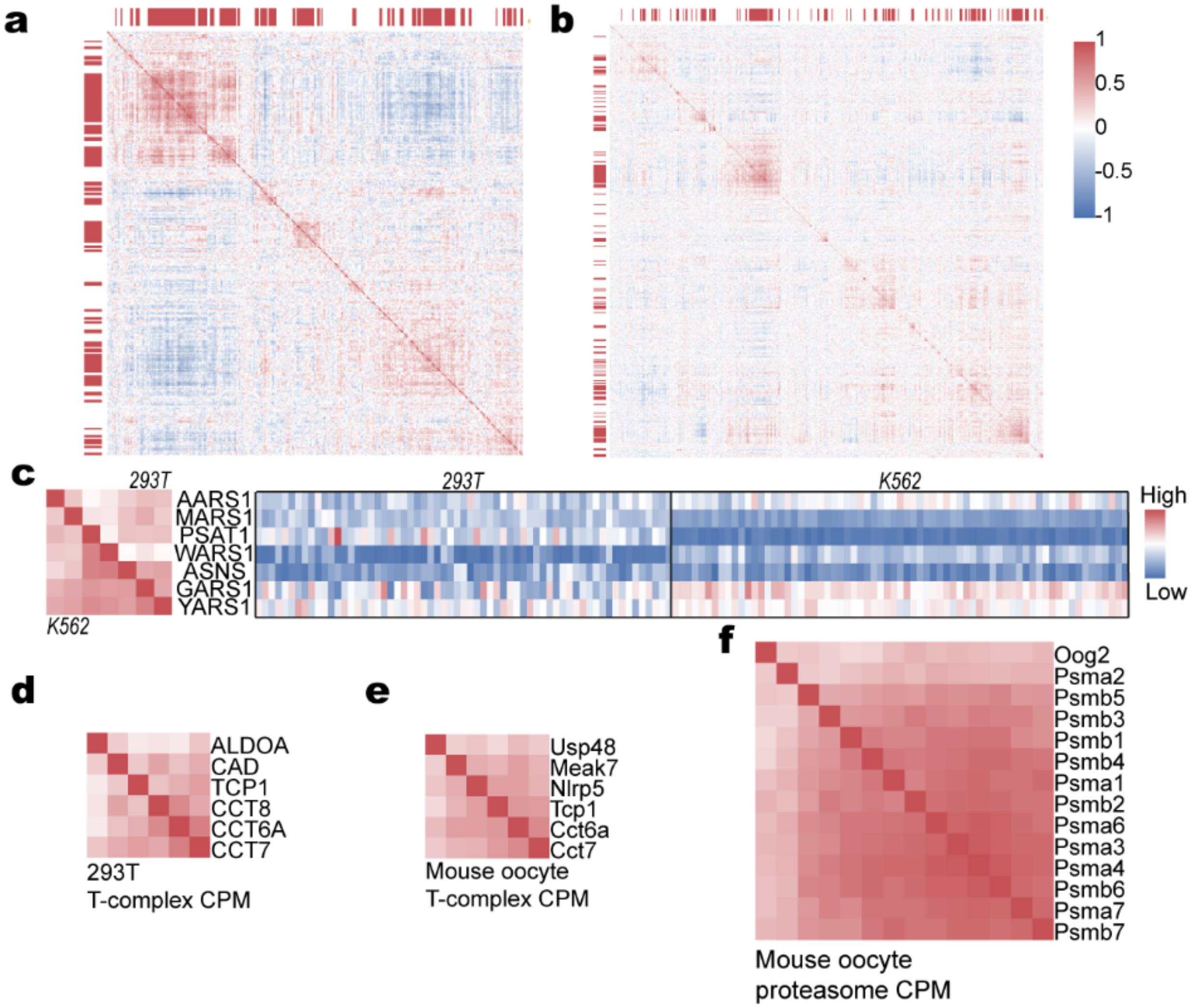


**Fig. S4.** Correlated protein modules are universal and cell-type specific. a. Correlation map of 1,150 proteins in 63 293T single cells. b. Correlation map of 3,422 proteins in 137 mouse oocytes. c. Comparison of tRNA aminoacylation modules of K562 and 293T. Proteins not quantified in both cell lines were removed. Heatmap represents the relative abundances of corresponding proteins in different single cells of K562 and 293T. d. T-complex CPM of 293T. e. T-complex CPM of mouse oocyte. f. Proteasome CPM of mouse oocyte.
